# Deciphering Metabolic Pathways and Protein-Protein Interaction Networks in Ankylosing Spondylitis through Single-Cell RNA Sequencing

**DOI:** 10.1101/2024.10.21.619465

**Authors:** Merve Yarıcı, Muhammed Erkan Karabekmez

## Abstract

Ankylosing Spondylitis (AS) is a common autoimmune disease affecting spinal joints and causing chronic pain. Understanding the roles of different cell types in AS can facilitate the development of effective treatments. In this study, we analyzed scRNA-seq data of peripheral blood mononuclear cells (PBMC) from AS patients and healthy controls collected from the literature. Using the GIMME algorithm, we created genome-scale metabolic models for each cell type to analyze reaction fluxes varying between patient and healthy conditions. Our findings revealed increased purine metabolism flux, fatty acid degradation, and glycolysis in CD14 monocytes, CD4 memory, CD4 naive, and CD8 T cells in AS patients compared to healthy individuals. Additionally, by integrating multi-omics approaches we generated cell-type-specific protein-protein interaction (PPI) networks, uncovering 63 rewired hubs across nine cell types. RPS11 emerged as the most significant hub, essential in translation and there are evidences in the literature that implicate it in AS. These results provide a detailed understanding of the metabolic and protein interaction changes in specific immune cell types in AS, highlighting RPS11 as a critical regulatory hub that could serve as a potential biomarker or therapeutic target for developing more precise and effective treatments.

## 1. Introduction

The human body contains numerous complex systems, with the immune system being one of the most crucial. The immune system protects against diseases, preventing or delaying the proliferation and spread of pathogens and tumor cells. It protects its own tissues and molecules while combating foreign invaders [1][2].In some instances, the immune system cannot distinguish between the body’s own healthy cells and foreign cells or molecules, leading to an attack on its own cells. This dysregulation in immune recognition and response results in the development of autoimmune diseases, where the body mistakenly targets its own cells, leading to chronic inflammation and tissue damage. While some autoimmune diseases, such as Type 1 diabetes, multiple sclerosis, lupus, and psoriasis, are relatively easy to diagnose, many others have pathogenesis that remains unclear, making early detection and treatment more challenging [2].

Ankylosing Spondylitis (AS) is one of the most common autoimmune diseases and a prevalent type of Spondyloarthritis (SPA), a broad group of inflammatory rheumatic diseases [3]. AS primarily affects the spinal joints, leading to chronic pain and reduced quality of life [4]. Multiple immunological, genetic, and non-genetic risk factors contribute to the onset of AS, but its pathogenesis is still unclear. Therefore, identifying biomarkers for early diagnosis is crucial. While the HLA-B27 gene is strongly associated with AS, it is not a reliable biomarker due to a high rate of false positives [6]. Additionally, there is no definitive cure for AS; current treatments focus on slowing disease progression and alleviating symptoms [7].

Integrating omics technologies like transcriptomics, methylomics and interactomics in AS studies provides an opportunity to uncover molecular mechanisms underlying the disease and identify potential drug targets and new biomarker candidates [8][9]. Multi-omics approaches offer the ability to analyze complex biological systems by integrating various biological data with a more holistic view of biological processes and interactions within living organisms [10]. In autoimmune diseases, Peripheral Blood Mononuclear Cells (PBMCs) are often used as they provide a rich source of immune-related gene expression data. Through experimental methods like qRT-PCR, key genes with significant expression changes can be identified and measured. Computational methods, such as bulk RNA-seq and single-cell RNA-seq (scRNA-seq), can be employed to relate these findings by providing a deeper, transcriptome-wide perspective on gene expression [11] [12].

Understanding the roles of different cell types in the treatment development process is important for complex structures including tumors or autoimmune diseases with heterogeneous cell compositions, whose pathogenesis remains unclear due to immunological and genetic factors. Analyzing cells individually with the single-cell RNA sequencing (scRNA-seq) method makes it possible to characterize cells associated with the disease and identify distinct cell types [13].

Immune cells provide the primary response to abnormal situations, making it crucial to examine each cell type in its biological context to understand their specific roles and immunological responses to diseases [14]. Different immune cell types, specifically NK cells, CD4+ T cells, and Th17 cells, play a role in the pathogenesis of AS [4]. Therefore, the single-cell RNA-seq (scRNA-seq) technique is crucial for offering a more precise understanding of their roles in AS.

Metabolism plays a pivotal role in immune cell activation and function, making it highly relevant in the study of autoimmune diseases including AS. Immune cells have distinct metabolic demands that support their activation, differentiation, and function. Targeting these metabolic pathways provides a more selective approach to regulating immune responses, potentially offering therapeutic advantages over broad immunosuppression [15]. Recent studies have shown that metabolic reprogramming can modulate the balance between effector and regulatory immune cells, which could be key in controlling autoimmunity [16][17].

Integrative approaches in system biology help to understand the complex structures and metabolic functions of organisms [18]. In this context, Genome-scale metabolic models (GSMMs) emerged as powerful frameworks for simulating and analyzing the metabolic networks of organisms at a genome-wide scale. GSMMs are mathematical simulations of an organism’s metabolism. This creates a framework for applying systems biology approaches across various fields, from medicine to biotechnology [19] [20]. GSMMs, which include all known metabolic genes and reactions in an organism, are primarily used to gain insights into the transcriptional regulation of metabolism, understand intracellular functioning, examine conditions that cause metabolic disorders, and investigate disease mechanisms in detail [21]. Additionally, GSMMs can provide faster results than experimental methods for identifying gene-reaction associations and predicting metabolic fluxes [22].

Moreover, biological networks characterize biological processes and provide essential clues to understanding the complex mechanisms inside and outside the cell [23]. To investigate the immunological mechanisms of autoimmune diseases or heterogeneous diseases understanding the interaction networks within and between cells is an important step in developing effective treatment methods. There are more than 22,000 protein-coding genes in the human genome, and the interactions of these proteins with each other and with other molecules mediate many biological processes [24] [25]. Multi-omics approaches, which integrate diverse datasets such as the interactome, transcriptome and are increasingly employed to study complex biological systems [8]. Protein-protein interaction (PPI) networks help make sense of the complex interactions within and between cells [26] . In biological systems, analyzing protein interactions and identifying key nodes (hubs) within PPI networks help reveal the underlying mechanisms and roles of proteins in complex processes [27].Therefore, finding topologically altered regions in a PPI network can give clues about biological switches across different contexts. Network comparison and identification of rewired hubs in which topology changes the most in different networks can help unveil biological mechanisms [28]. Network topologies differ in different cell types; it is important to identify rewired hubs by comparing cell type-specific networks to study the immunological mechanism of a heterogeneous disease or to investigate individual-specific disease states [29].

In this study, scRNA-seq datasets of AS were analyzed to elucidate the biological mechanisms of the disease and to identify potential biomarkers for treatment development. Cell type-specific metabolic models were constructed to create mathematical simulations of metabolism. Additionally, cell-type-specific PPI networks were created to understand the interaction networks within and between cells. This study also proposed an approach for discovering new biomarkers for complex biological processes such as autoimmune diseases and cancer.

## 2. Materials and Methods

### 2.1. Selection of scRNA-seq Datasets

To investigate AS mechanisms, PBMC scRNA-seq datasets were used because they contain key immune cells, which play a crucial role in responding to disease and abnormal conditions. The datasets taken from the GEO database [30] via GSE163314 [31] and GSE194315 [4] accession codes. Transcriptomic data from PBMC samples of 2 healthy individuals and two individuals with axSpA, a subtype of AS, in the dataset with access code GSE163314 (DS1) and additionally, from 10 individuals with AS and 29 healthy individuals in the dataset with access code GSE194315 (DS2) were used.

### 2.2. Analysis of scRNA-seq Data Sets and Detection of Differentially Expressed Genes

scRNA-seq data analysis of the healthy and patient groups for both data sets was performed by downloading the “matrix”, “features” and “barcodes” files from the GEO database and using the Seurat 4.3.0 package [32] on the R 4.2.3 platform. Quality control for transcriptomic data of PBMC samples for each individual in both datasets was performed using commonly applied metrics [33] to filter out cells with more than 4000 or less than 200 unique genes, cells with a total number of molecules less than 20000, and cells with a mitochondrial RNA percentage greater than 5%.

Seurat objects of all AS patients and all healthy controls in both data sets were combined to form a single healthy and AS patient object in each data set. These two Seurat objects, containing transcriptomic data from PBMC samples of healthy controls and AS patients, were then merged into one seurat object for further analysis.

To determine the size of the combined Seurat Objects for DS1 and DS2, cells were clustered according to PCA (principal component analysis) scores, and 20 PC (principal component) were selected for both data sets. Also, to correct the batch effect, the “FindIntegrationAnchors” function was used to find cells representing the same fundamental biological entity as “anchors” among the cells in different data sets in the same Seurat object, and data integration was performed with the IntegrateData function.

In DS1, data integration was performed with dimensions set to 1:20 and resolution set to 0.1, while in DS2, dimensions were set to 1:20 and resolution to 0.6. After clustering both datasets, the Wilcoxon rank sum test [34] was applied, and each cluster was compared with the other clusters one by one. Marker genes were identified, and the cell types corresponding to each cluster were determined. Subsequently, differentially expressed genes (DEGs) between the healthy control group and AS patient data were identified for each cell type with non parametric Wilcoxon rank sum test. Genes with an adjusted p-value of less than 0.05 were selected as DEGs.

### 2.3. Construction of Cell Type-Specific Genome-Scale Metabolic Models

For each group in DS1 and DS2 (AS patients and healthy controls), separate genome-scale metabolic models (GSMMs) specific to each cell type were constructed using the GIMME algorithm [15] of Cobra Toolbox [35] was used by utilizing MATLAB 2023 platform [36]. Before using the GIMME algorithm, log normalization was applied to the gene expression values of each cell type, and genes with zero total expression were removed. The gene lists of each cell type were then matched with Recon3D [37], and only the expression data of the genes included in Recon3D were used. The ‘mapExpressionToReaction’ function in the Cobra Toolbox [35] was used to map gene expression data to their corresponding metabolic reactions.

For each dataset, thresholds for the GIMME algorithm were determined by averaging the expression values of each gene across all samples for each cell type, and identifying the median value of them.

Flux balance analysis (FBA) [38] was performed to analyze the fluxes of biochemical reactions within the metabolic network and to predict ATP- demand in AS and healthy-control specific genome-scale metabolic models (GSMMs) created for each cell type. For the FBA, the Cobra Toolbox 2.39.0 [35] and the Gurobi optimization package were utilized on the Matlab platform [36].

### 2.4. Flux Sampling and Reaction Set Enrichment Analyses

The Cobra Toolbox [35] software was used for flux sampling. The ACHR (artificial centered hit-and-run) algorithm [39] was employed as a sampler using the sampleCbModel function, and 2000 flux points were sampled for each reaction within each GSMM. Flux sampling was performed separately for both conditions (patient and healthy) in each cell type. The two-sample Kolmogorov-Smirnov test [40] was applied using the kstest2 function in Matlab to detect fluxes varying between the two conditions. Additionally, pairwise distances of sampling distributions between the two conditions were calculated using the Wasserstein distance [41], a metric that quantifies the distance between two probability distributions, in Matlab. The Wasserstein distance was previously used in the analysis of GSMMs to compare metabolic flux distributions [42]. To normalize the distance values, the interquartile range of each reaction for each sample was calculated, and the fold change was determined as the ratio of the Wasserstein distance to the interquartile range. Reactions with a corrected p-value below 0.01 and a fold change value above one were identified as differentially expressed reactions. For each reaction, significantly increasing (Differentially Up Flux, DUF) and decreasing (Differentially Down Flux, DDF) fluxes were determined by subtracting the average flux value in the healthy condition from the average in the diseased condition. Reactions with a negative mean value were classified as DDF, while reactions with a positive mean value were classified as DUF.

In both datasets, enrichment analyses of DUFs and DDFs determined for each cell type were performed using the RSEA software tool [43].

### 2.5. Construction of Context-Specific Protein-Protein Interaction Networks for Each Cell Type

For the DS2, the cell types with a sufficiently high number of DEGs were selected for further PPI network construction. For this purpose, the cell types that brought less than 100 DEGs were excluded from the rest of the study.

The Sørensen–Dice Coefficient (SDC) [44] was used to calculate the similarity between cell-type-specific DEG lists. While calculating this coefficient, the shared DEGs of the two cell types were multiplied by two and divided by the sum of the DEG numbers of the cell types (Equation 1).

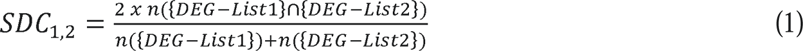

The PPI networks for the proteins encoded by DEGs identified for specific cell types with more than 100 DEGs were created using the human protein interaction dataset downloaded from the String database [45]. First, protein interactions detected for each cell type were visualized using the Cytoscape 3.9.0 [46] platform, and topological investigations of the networks were performed. Then, by using the DyNet [47] application of Cytoscape through pairwise combinations of cell types, the rewired hubs with the most differentiation in the topological role were determined according to the DyNet Rewiring Score (Dn-score). The top ten hubs with the highest Dn-scores were identified as rewired hubs for each cell type combination. The rewired hubs that changed consistently across multiple cell type comparisons were identified as key differential hubs.

Enrichment analysis of key differential hubs was performed and visualized using the enrichplot R package [48] on the R 4.2.3 platform.

## 3. Results

The scRNA-seq data analyses were conducted separately for each dataset using a single Seurat object, containing 22,440 cells in DS1 and 47,414 in DS2.

For each dataset, similar cells were clustered and visualized using UMAP. Twelve clusters were created in DS1 and 22 in DS2 (Fig 1).

**Fig. 1.**
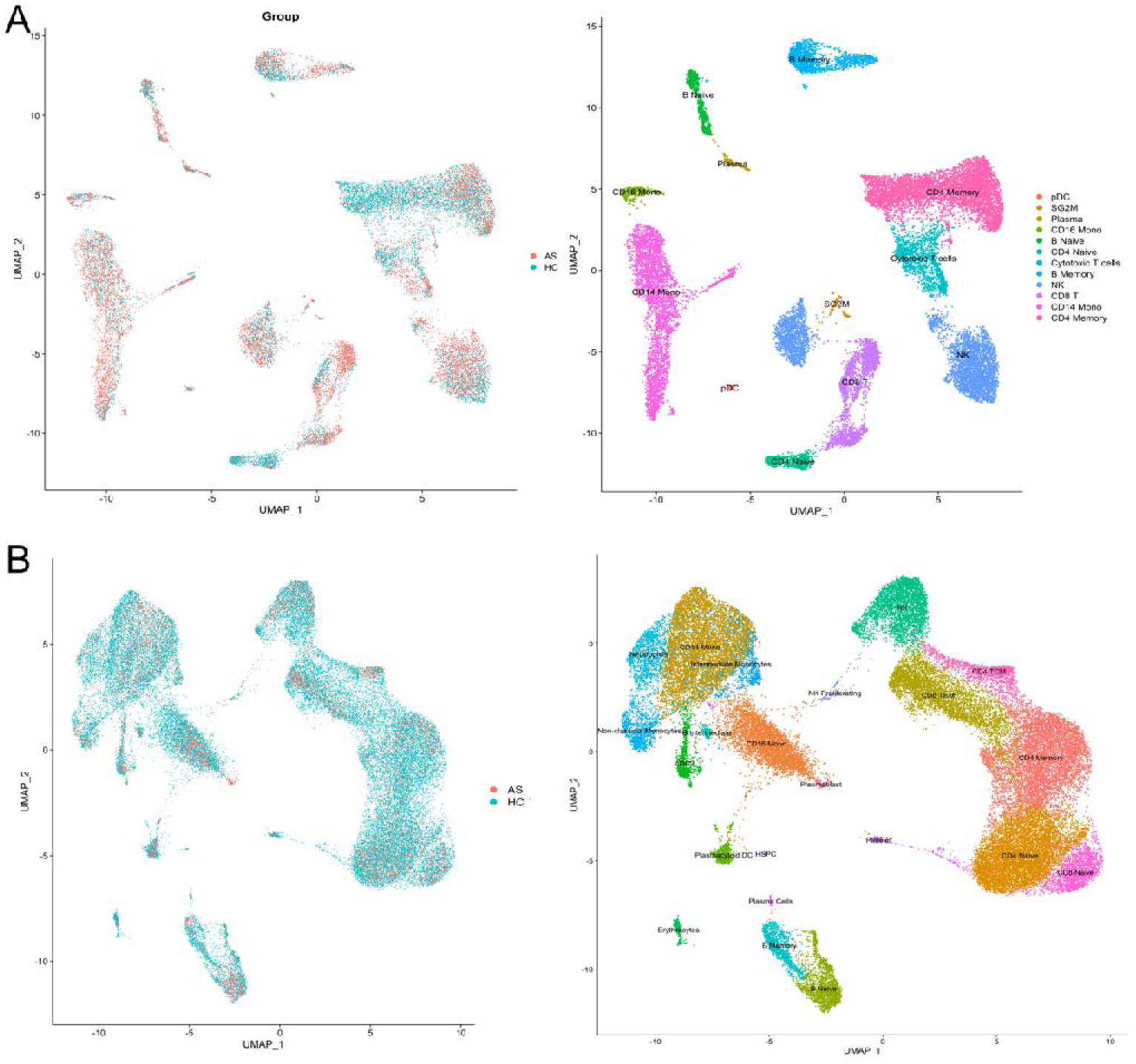
UMAP visualizations for DS1 (A) and DS2 (B): The left graphics show the distribution of cells between the diseased and healthy conditions, while the right graphics show the distribution of the identified cell clusters.

After the clustering process, the differentially up-regulated genes were examined to determine which cluster represented each cell type. The Wilcoxon rank-sum test was applied to detect these genes. By comparing each cluster with other cluster groups, differentially expressed genes (DEGs) were identified for each cluster (Table S1). In DS1, clusters 2 and 4 represented the same cell type based on DEG analyses, and the analyses continued with 12 distinct cell types (Fig.1).

For the cell types detected in each dataset, genes with differentially expressed levels between the diseased and healthy conditions were identified, and those with a corrected p-value below 0.05 were considered DEGs (Table 1).

**Table 1.** DEG numbers for cell types identified in DS1 and DS2.

| Cell Types | DS1- GSE163314 |  |  | DS2- GSE194315 |  |  |
| --- | --- | --- | --- | --- | --- | --- |
|  | DEG | Up-regulated DEGs | Down-regulated DEGs | DEG | Up-regulated DEGs | Down-regulated DEGs |
| CD14 Mono | 411 | 245 | 166 | 461 | 344 | 117 |
| CD16 Mono | 55 | 25 | 30 | 3 | 2 | 1 |
| CD4 Memory | 636 | 447 | 189 | 342 | 277 | 65 |
| CD4 Naïve | 51 | 32 | 19 | 406 | 329 | 77 |
| B Naïve | 86 | 47 | 39 | 158 | 110 | 48 |
| B Memory | 184 | 88 | 96 | 24 | 6 | 18 |
| NK | 245 | 145 | 100 | 149 | 95 | 54 |
| Plasma | 0 | 0 | 0 | 1 | 0 | 1 |
| Plasmacytoid DC | 11 | 7 | 4 | 26 | 9 | 17 |
| CD8 T | 264 | 160 | 104 | - | - | - |
| SG2M | 2 | 0 | 2 | - | - | - |
| Cytotoxic T cells | 171 | 82 | 89 | - | - | - |
| B Intermediate | - | - | - | 9 | 1 | 8 |
| CD4 TCM | - | - | - | 115 | 54 | 61 |
| CD8 NAIVE | - | - | - | 156 | 103 | 53 |
| CD8 TEM | - | - | - | 200 | 124 | 76 |
| cDC2 | - | - | - | 131 | 69 | 62 |
| Intermediate Monocytes | - | - | - | 49 | 26 | 23 |
| NK Proliferating | - | - | - | 1 | 0 | 1 |
| Non-Classical Mono | - | - | - | 33 | 13 | 20 |
| Plasmablast | - | - | - | 63 | 0 | 63 |
| Platelet | - | - | - | 12 | 0 | 12 |

Genome-scale metabolic models specific to each cell type were constructed using the GIMME algorithm. The threshold values were 0.06 for DS1 and 0.08 for DS2. The objective function was set as the demand for ATP (’DM_atp_c_’). As a result of FBA applied to each cell type-specific GSMM, significant differences in ATP demand were observed between AS patients and healthy controls (HC). Overall, specific cell types, such as B Intermediate cells and cDC2 cells, in AS patients showed increased ATP demand compared to healthy controls, indicating heightened metabolic activity in these diseased states. Conversely, other cell types, like B Memory cells and CD4 Memory cells, exhibited decreased ATP demand in AS patients, reflecting reduced metabolic activity. These variations demonstrate that ankylosing spondylitis affects the metabolic demands of different PBMC cell types in distinct ways, highlighting the disease’s complex impact on cellular metabolism (Table S2, Table S3).

After flux sampling, DERs were determined for each dataset. For example, in the DS1, CD14 Monocytes exhibited the highest number of up-regulated DERs (989) and down-regulated DERs (475). Similarly, in the DS2, B Intermediate cells showed significant regulation with 321 up-regulated and 286 down-regulated DERs. Other notable findings include up-regulated and down-regulated DERs in CD16 Monocytes, CD4 Memory, CD4 Naive, and CD8 T Cells, among others (Table 2). Additionally, pathway enrichment analyses were performed using the RSEA software tool to differentially increase (DUF) and decrease fluxes (DDF) between healthy and diseased conditions in each cell type.

**Table 2.** Numbers of DUF and DDF in DS1 and DS2.

| DS1- GSE163314 |  |  | DS2- GSE194315 |  |  |
| --- | --- | --- | --- | --- | --- |
| Cell Type | DUF | DDF | Cell Type | DUF | DDF |
| CD14 Mono | 989 | 475 | B Intermediate | 321 | 286 |
| CD16 Mono | 343 | 351 | B Memory | 141 | 117 |
| CD4 Memory | 258 | 215 | B Naïve | 69 | 51 |
| CD4 Naïve | 257 | 196 | CD4 Memory | 223 | 154 |
| CD8 T Cells | 282 | 278 | CD4 Naive | 159 | 151 |
| Cytotoxic T cells | 339 | 303 | CD4 TCM | 259 | 175 |
| B Naïve | 79 | 59 | CD8 NAIVE | 137 | 85 |
| B Memory | 219 | 178 | CD8 TEM | 50 | 36 |
| NK | 420 | 296 | CD14 Mono | 207 | 136 |
| Plasma | 19 | 14 | CD16 Mono | 72 | 56 |
| SG2M | 331 | 251 | cDC2 | 377 | 327 |
| PDC | 734 | 559 | Erythrocytes | 34 | 20 |
|  |  |  | HSPC | 533 | 303 |
|  |  |  | Intermediate Monocytes | 454 | 348 |
|  |  |  | Neutrophils | 194 | 143 |
|  |  |  | NK | 102 | 80 |
|  |  |  | NK Proliferating | 589 | 403 |
|  |  |  | Non-Classical Mono | 287 | 183 |
|  |  |  | Plasmablast | 0 | 68 |
|  |  |  | Plasma cells | 0 | 76 |
|  |  |  | Plasmacytoid DC | 317 | 256 |
|  |  |  | Platelet | 43 | 50 |

Additionally, pathway enrichment analyses were performed using the RSEA software tool for differential up (DUF) and down fluxes (DDF) between healthy and diseased conditions in each cell type. Overall, fluxes were increased in purine metabolism, fatty acid degradation, and glycolysis in CD14 Monocytes, CD4 Memory, CD4 Naive, and CD8 T cells of AS patients compared to healthy individuals (Table S4, Table S5).

For the construction of PPI networks for each cell type in DS2, nine cell types were found to have more than 100 DEGs. These nine cell types are: CD14 Mono, CD4 Memory, CD4 Naïve, B Naïve, NK, CD4 TCM, CD8 Naïve, CD8 TEM, and cDC2 (Table 1).

The PPI networks created for binary combinations of nine cell types created by DyNet are shown in the upper triangle of Fig. 2. The numbers of remaining nodes in the networks are shown in the diagonal of the figure. In the lower triangle of the figure, the number of aligned nodes of each combination of the cell types and the similarity coefficient of the cell-specific DEG lists calculated by using the Sørensen–Dice coefficient statistics are tabulated (Fig.2). The highest similarity was observed between CD8 TEM and NK cells and the most distinct DEG list was found in CD14 mono cells. The low similarity of CD4 Monocytes DEGs is partly because of the size dependence of SDC, as this cell type has the largest DEG list.

**Fig. 2.**
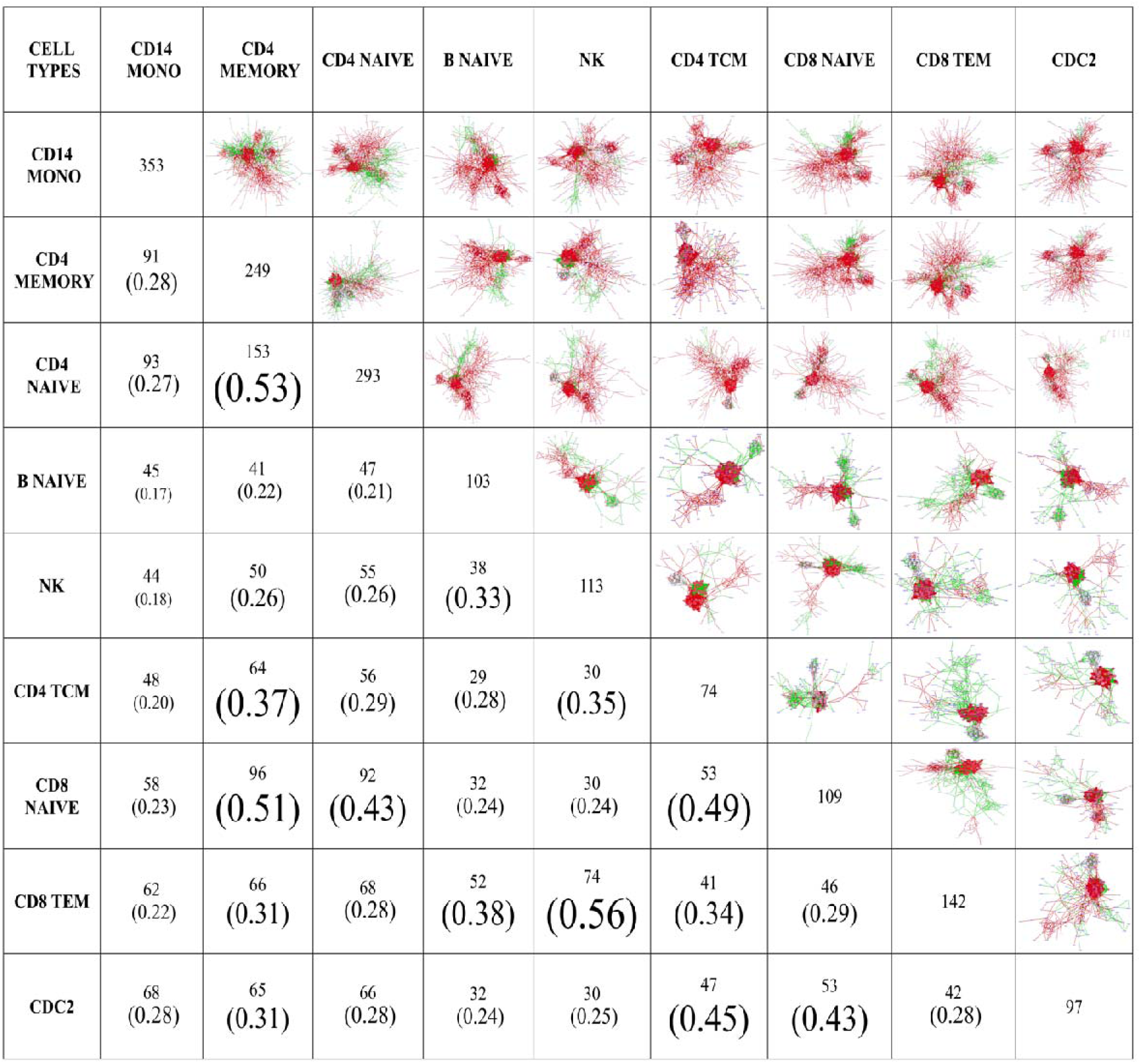
Visualization of aligned PPI networks of seven cell types with DyNet: The network images of binary combinations of cell types (upper triangle) and the number of nodes in the combination (lower triangle) and number of nodes (common diagonal) are shown in the table. Sørensen–Dice coefficients of DEG lists are indicated in parentheses for each cell type combination with a font proportional to the values.

The top ten most rewired hubs with the highest Dn scores for each cell type pair were identified, resulting in a cumulative total of 63 hubs. Among these, fifteen hubs that appeared in at least ten cell-type pairs were arbitrarily designated as key differential hubs (KDH). These hubs are: RPS11, RPL13A, RPS28, RPS3A, RPS10, RPS27A, RPL11, RPS8, EEF1A, RPS9, RPL9, RPS12, UBA52, RPS15A, and RPS4X. RPS11, a ribosomal protein that contributes to the structure and function of ribosomes and regulates protein synthesis, emerged as the most consistently rewired KDH, significantly altered in 28 combinations. The variations in the intersection of RPS11 neighbors across cell type-specific PPI network combinations can be attributed to the distinct regulatory mechanisms and functional roles of RPS11 in different cell types (Fig.3). The enriched biological processes associated with the KDHs are predominantly related to cytoplasmic translation, ribonucleoprotein complex biogenesis, and rRNA processing, which highlights the central role of ribosomal proteins in protein synthesis and cellular metabolism. The most enriched process, cytoplasmic translation, is linked to RPS11’s function in facilitating the formation of ribosomes and contributing to translation [49] (Fig. 4).

**Fig 3.**
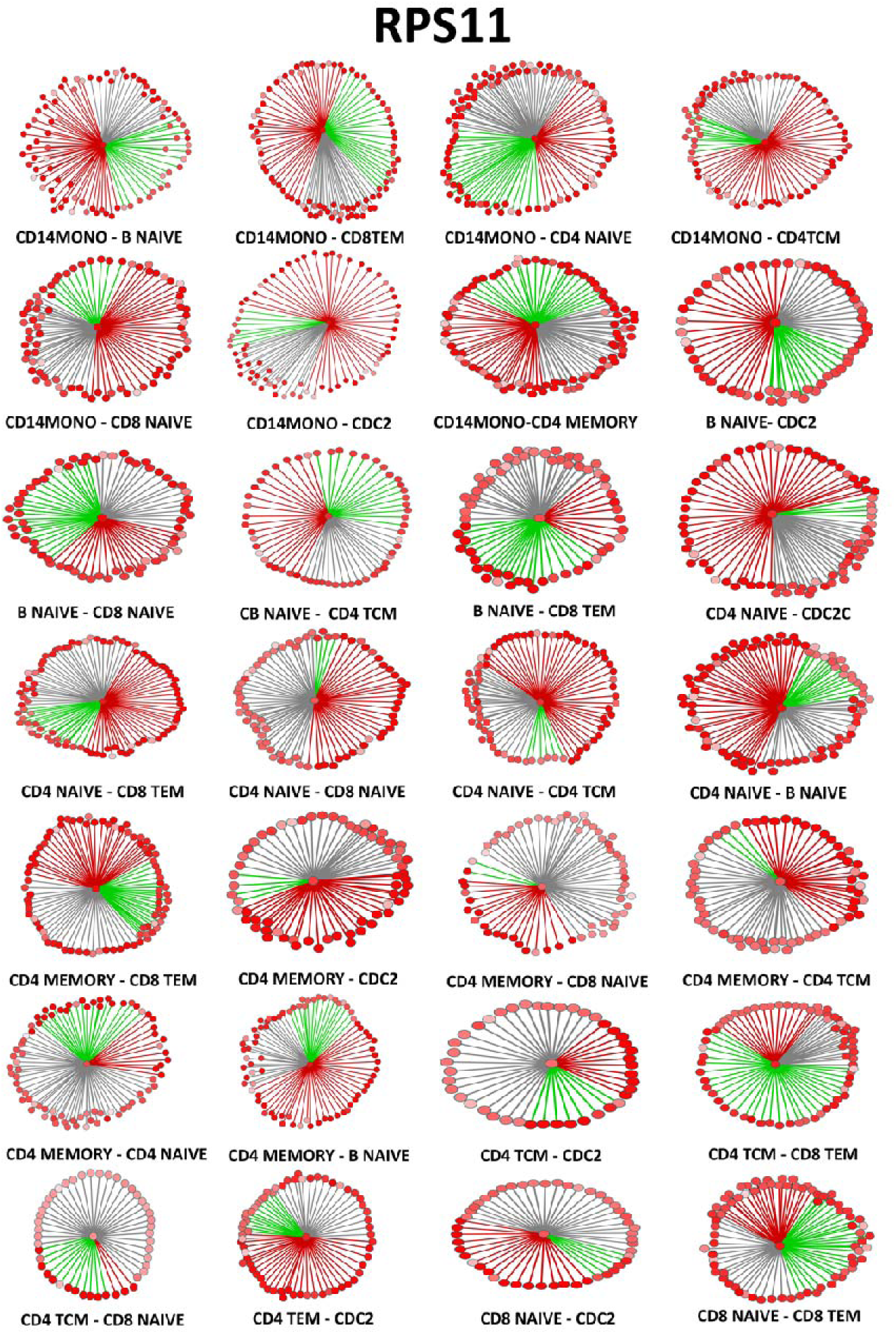
Change in neighbors of RPS11 combinations of cell types. In pairwise combination networks, the first specified cell type is represented by red edges, while green edges represent the second cell type. The gray edges represent common edges that are included in both cell-type networks. Node colors are highlighted according to the DyNet Rewiring Score (Dn- score), with a higher proportion of red indicating a greater score of rewiring.

**Fig. 4.**
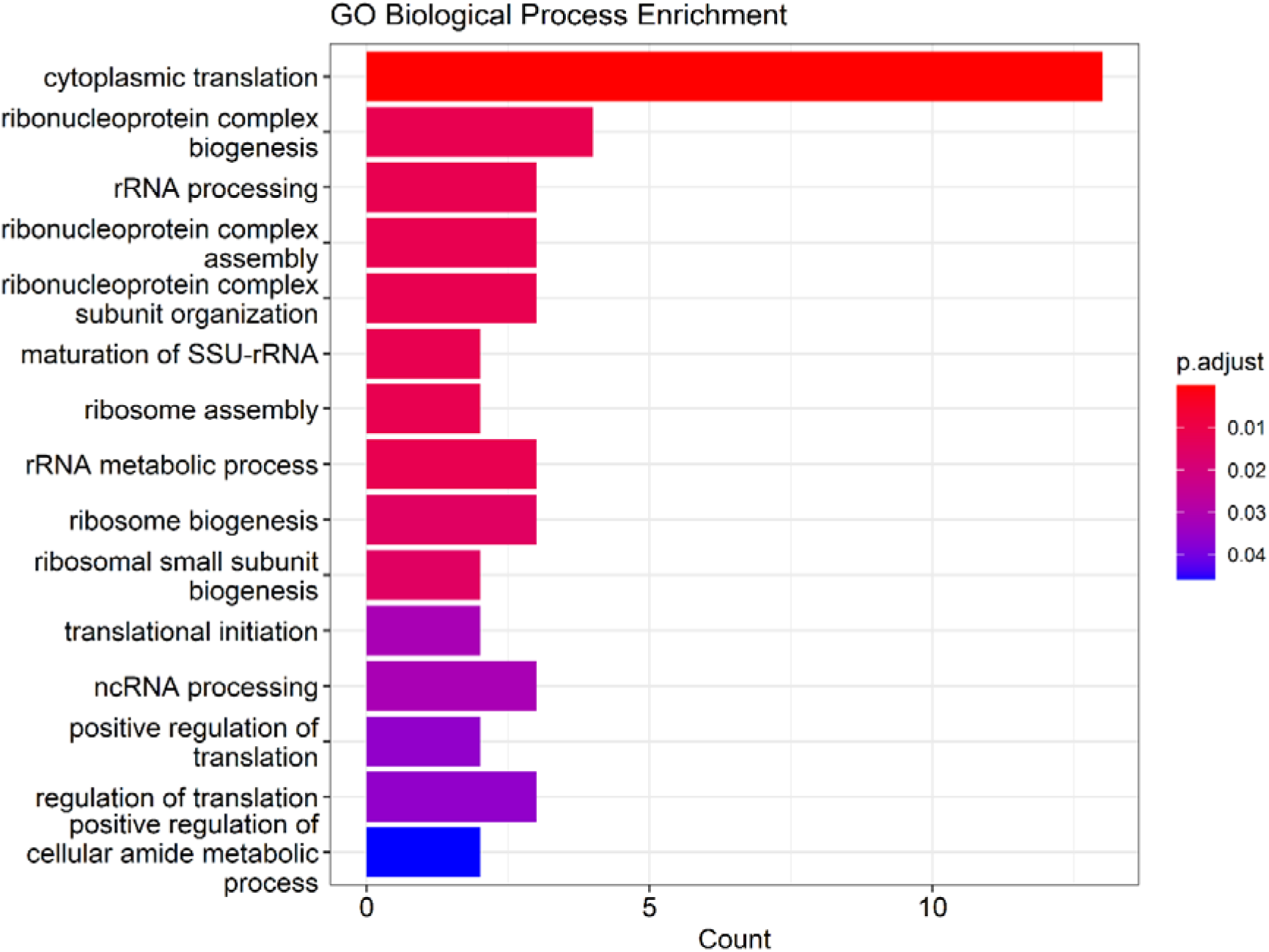
E**n**richment **Analysis of KDHs** Gene Ontology (GO) Biological Process Enrichment analysis conducted for the identified key differential hubs (KDHs), particularly highlighting the top enriched biological processes. The color-coded bars represent the significance levels (adjusted p-values) of the enriched terms, with the most significant processes at the top.

## 4. Discussion

Investigating the immunological mechanisms of autoimmune diseases, such as ankylosing spondylitis (AS), involves understanding the roles of various cell types in biological processes, facilitating effective treatment development. In this study, AS PBMC scRNA-seq data were analyzed using Seurat, and genome-scale metabolic models specific to each cell type were created using the GIMME algorithm. In the metabolic models created, differentially increasing and decreasing fluxes were detected between patients with AS and the healthy control group.

Understanding metabolism in the context of autoimmune diseases is particularly relevant because immune cell activation, proliferation, and differentiation are linked to metabolic pathways [50]. Metabolic mechanisms are the hallmark of immune response, and changes in energy production, biosynthesis, and immune signaling contribute directly to disease pathogenesis [16]. In autoimmune diseases like AS, altered metabolic activity in immune cells can fuel chronic inflammation and tissue damage, underscoring the importance of studying metabolism in such conditions.

In DS1, 12 clusters were detected using UMAP; however, DEG analysis revealed that clusters 2 and 4 both represented NK cells and were thus considered a single cluster.. This variation in UMAP positioning may result from data distortion in two-dimensional reduction, a known issue supported by other studies [51]. As a result of scRNA-seq data analysis, four cell types with more than 200 DEGs were detected in both datasets, including CD14 Monocytes, which play a crucial role in AS pathogenesis [52] (Table 1). These cells have been found to be increased in the peripheral blood of AS patients compared to healthy individuals [53] [54]. Consequently, it was concluded that CD14 Monocytes contribute to the inflammatory response and pathogenesis of AS.

A two-sample Kolmogorov-Smirnov test [31] was applied to identify differing fluxes between diseased and healthy conditions for each reaction in every cell type, as the flux samples were not normally distributed. However, the p-values did not provide a robust filtering criterion.

Genome-scale metabolic models specific to CD14 Monocytes for AS and healthy control conditions revealed that purine and pyrimidine metabolism fluxes were differentially increased in both datasets potentially exacerbating tissue damage [55], while TCA cycle fluxes were reduced, likely due to inflammation-induced metabolic shifts [56] (Table S4, Table S5). This suggests changes in metabolic activity and energy production pathways in CD14 Monocyte cells, as the TCA cycle is a crucial metabolic pathway involved in energy production and various cellular processes.

Additionally, the flux in propanoate metabolism, which involves the utilization, synthesis, and degradation of propanoate—a fatty acid important for cellular metabolism—was differentially downregulated in AS patients compared to healthy controls.Propanoate is initially produced in the intestine through microbial fermentation as a byproduct of intestinal microbiota metabolic activities. Considering the changes in gut microbiota composition and function in AS patients, the production and availability of short-chain fatty acids like propanoate may be affected. Alterations in the gut microbiota of AS patients could lead to reduced propanoate availability [57] [58].

Additionally, CD4 Memory, CD4 Naive, and CD8 T cells undergo less apoptosis in AS patients than in healthy individuals, resulting in higher prevalence in individuals with AS [59]. Studies have shown that memory T cells residing in the synovial fluid, peripheral blood, and intestines of AS patients are more abundant than in healthy individuals, with many of these cells recirculating to peripheral blood and inflamed joints [60] [61].

Our metabolic models for these cell types revealed that fluxes in purine metabolism, biosynthesis of unsaturated fatty acids, and glycolysis are differentially increased in CD4 Memory, CD4 Naive, CD8 T cells, and CD14 Monocytes in AS patients compared to healthy individuals. Additionally, fluxes in the arachidonic acid synthesis pathway, an unsaturated fatty acid, were up-regulated in CD4 Naive and CD8 TEM cells of AS patients. Previous studies have reported that differences in arachidonic acid metabolism in individuals with AS may contribute to inflammatory processes and disease pathogenesis [62] [63] [64]. Sundström et al. (2012) found a positive correlation between arachidonic acid levels in the plasma phospholipids of AS patients and the BASDAI index, which measures disease activity [63]. Therefore, targeting specific enzymes or receptors in the arachidonic acid pathway has the potential to be a therapeutic approach, given its contribution to AS pathogenesis.

Cell type-specific PPI networks revealed that ribosomal genes, particularly RPS11, which contribute to the structure and function of ribosomes and the regulation of protein synthesis, were significantly rewired across multiple cell types, highlighting their involvement in AS pathogenesis (Fig. 3). Ribosomal proteins like RPS11 may indirectly contribute to AS through immune dysregulation [65], aligning with similar findings in other autoimmune diseases [55].

Differences in the intersection of RPS11 neighbors in cell type-specific PPI networks reflect distinct regulatory mechanisms in different cell types. In AS, CD14 Monocytes play a key role in inflammation, producing pro-inflammatory cytokines like TNF, IL-23, and CCL17, contributing to chronic inflammation and joint damage [52] [67]. CD14 Monocytes also have enhanced antigen presentation capacity and activate CD8 T cells, further contributing to AS pathogenesis. The minimal intersection of RPS11 neighbors between CD14 Monocytes and CD8 TEM cells suggests cell type-specific protein interactions. Similarly, the small intersection in CD14 TCM cells indicates distinct molecular profiles, whereas CD4 Naive, CD4 Memory, and CD8 Naive cell networks show a larger intersection due to shared immune response mechanisms.

Beyond RPS11, a subset of ribosomal proteins was identified as key differential hubs (KDHs) through enrichment analysis, with notable associations with translation, ribosome biogenesis, and RNA processing (Fig. 4). Perturbed ribosomal protein expression has been linked to disorders such as cancer, ribosomopathies, and autoimmune diseases, including AS [65] [66]. Ribosomal genes may not directly cause AS but are likely affected by immune system dysregulation, indirectly influencing cellular processes. Further research is needed to understand the role of ribosomal genes like RPS11 in AS.

As a result, this study performed scRNA-seq data analysis for AS. Utilizing the scRNA-seq technique, a relatively new method studied only in recent years, genome-scale metabolic models were created for each detected cell type. Additionally, integrated analyses of protein interaction networks were conducted to investigate the biological mechanisms of AS and identify alternative targets for treatment development. This study also proposed an approach for discovering new biomarkers for complex biological processes including autoimmune diseases and cancer.

## Contributions

**Merve Yarıcı:** Conceptualization and methodology; data analysis and interpretation; writing original draft preparation; resources; validation .

**Muhammed Erkan Karabekmez:** Conceptualization and methodology; writing original draft preparation; writing proofreading; supervision.

